# Streamlined Montage Cryo-Electron Tomography for Exploring the Ultrastructure of Cells and Tissues

**DOI:** 10.1101/2025.09.01.673430

**Authors:** Ryan Hylton, Micaela Boiero Sanders, Adriana Prajica, Gavin Rice, Stefan Raunser

**Affiliations:** Department of Structural Biochemistry, Max Planck Institute of Molecular Physiology, Otto-Hahn-Strasse 11, 44227 Dortmund, Germany

## Abstract

Cryo-electron tomography reveals the architecture of biological specimens at molecular resolution, albeit with a limited field of view. Montage tomography overcomes this limitation, but existing strategies depend on complex dose distribution imaging schemes or hardware modifications which prevent widespread adoption of the method. To lower the barrier to entry, we developed a straightforward workflow that deviates minimally from conventional image acquisition and leverages our software MontageMaker to merge overlapping tilt series. The resulting montage tomograms are produced with high throughput, retain nanoscale detail despite the uneven accumulation of dose, and are easily acquired on a wide variety of biological samples. Most importantly, this approach can be implemented immediately by any lab with access to automated cryo-ET data collection.

## Main Text

Among the structural biology methods, cryo-electron tomography (cryo-ET) is uniquely positioned to characterize the ultrastructure of biological specimens in their near-native state. Although capable of determining protein structures to sub-nanometer resolution^1,2^, perhaps its greatest strength is in revealing mesoscale cellular architecture and placing those high-resolution structures in their biological context. This strength is due, in part, to the high magnification used during imaging, but this is also responsible for one of its major limitations: a severely limited field of view (FOV). This can be alleviated by montage tomography, a technique where multiple overlapping tilt series are acquired on a region of interest and computationally merged to generate a single “montage” tomogram^3–8^. Performing montage tomography requires at least two operations that lie outside of a typical cryo-ET workflow. First, overlapping tilt series must be collected in such a way that minimizes sample damage while retaining nanoscale information. Then, the tilt images need to be stitched into montages.

Regarding tilt series acquisition: historically, montage tomography has mostly been performed on resin-embedded samples, owing to the radiation sensitivity of cryo-preserved ones and the apparent danger of accumulating dose on the overlapping regions^3–5^. There are only two examples in the literature where montage tomography is applied to vitrified samples which utilize complex imaging schemes to avoid these dose “hotspots”^6,7^. An alternative approach was recently introduced where a square electron beam substantially reduced uneven dose accumulation^8,9^. Though innovative, this concept still does not completely eliminate tile overlap since 1) some degree of overlap is required to create montages without seams and 2) even perfect tiling with no overlap is only so with a flat sample. Unless circumvented by specialized imaging schemes^8^, the beam effectively elongates into neighboring tiles at high tilt angles, just as in conventional tilt series acquisition. More importantly, circular beams are a well-established standard integrated into existing instrumentation and adopting square beam apertures requires modifications to functioning, costly microscopes – a shift that many users are hesitant to make without extensive validation and industry support.

Despite the imaging scheme, current strategies for subsequent tile stitching rely on custom scripts dedicated to specific workflows^6–8^, making this step difficult to perform if the data was collected otherwise. Perhaps the most useful tool to date comes from the MPACT (montage parallel array cryo-tomography) workflow which includes a script called “BlendStitch”^7^ that capitalizes on existing infrastructure within the software package IMOD^10^ for merging tiles. BlendStitch, however, expects tilt series to be imaged and processed using the remainder of the MPACT pipeline as inputs, making it unusable for data collected outside of that workflow.

Overall, while the above approaches successfully produce montage tomograms, the technique remains challenging to perform and is consequently seldom used. We therefore sought to provide a simpler solution to those seeking larger FOVs from their tomography data, first by tackling the simplest problem: tile stitching. To this end, we developed MontageMaker, a user-friendly GUI application. MontageMaker is built upon the BlendStitch script from the MPACT workflow^7^ and uses IMOD^10^ for its core functions. Unlike BlendStitch, however, MontageMaker can merge overlapping tilt series independent of acquisition software or imaging scheme and is agnostic to input file names, image order, etc. Additionally, it can merge many montages in succession for high-throughput processing (**Fig. 1 and Supplementary Fig. 1**).

**Fig. 1:**
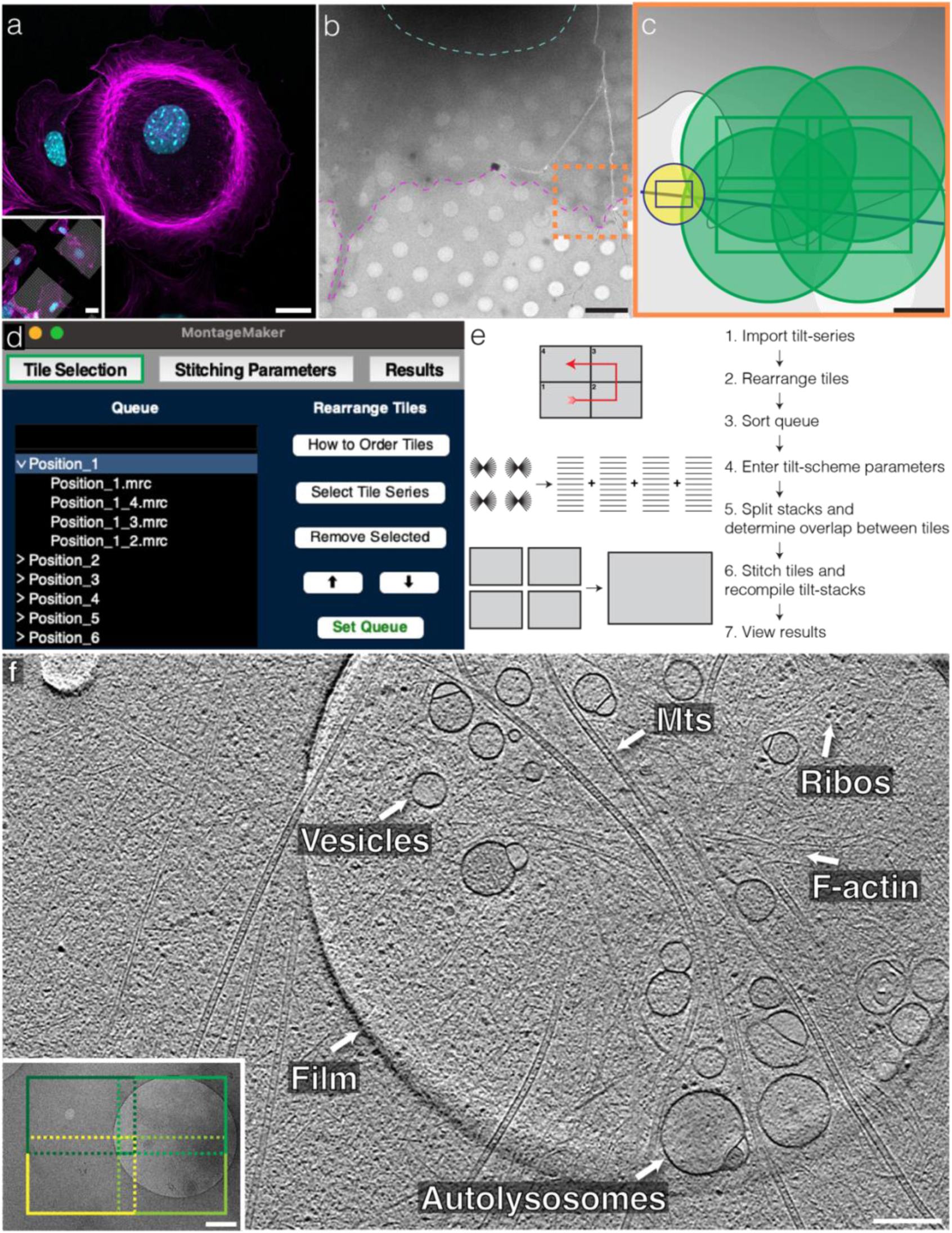
Streamlined montage cryo-electron tomography workflow. **a,** 10T1/2 fibroblasts growing on a glass slide (main) and an EM grid (inset). Magenta is actin (CellMask Deep Red Actin Stain) and cyan is the nucleus (NucBlue). Scale bars, 20 µm, **b,** TEM overview image showing a 10T1/2 cell on an EM grid. The cells spread enough to avoid sample thinning by FIB milling. The magenta line annotates the leading edge of the cell and the cyan one approximates the location of the nucleus. The orange dashed box shows the hypothetical size and location of the schematic in **c**. Scale bar, 5 µm. **c,** Cartoon representation of our scheme for overlapping tilt series collection in Tomography 5. Here, the focus and tracking positions (overlapping blue and yellow circles) are at a higher magnification than the acquisition sites (green) so as to limit the FOV lost to these activities. The blue line marks the tilt axis, rectangles show the location of the image on the camera, and the circles denote the size and location of the electron beam. Scale bar, 1 µm. **d,** Screenshot of the “Tile Selection” tab of MontageMaker. **e,** Overall workflow for importing and stitching tilt series into montages in MontageMaker. **f,** 7.2 nm-thick slice of a low-pass-filtered montage tomogram comprised of four individual tomograms with locations shown by the rectangles in the inset. Example microtubules (Mts), ribosomes (Ribos), filamentous actin (F-actin), vesicles, autolysosomes, and the edge of a hole in the grid’s film are labeled. Scale bars, 200 nm and 500 nm (inset).

With tile stitching streamlined, we concentrated on image acquisition. Previous methods aimed to keep dose accumulation as low as possible by spreading it evenly across the montages^6,7^. Such efforts may be unnecessary, however, given the primary purpose of montage tomography: large-scale overviews for ultrastructural analysis. Especially given the absence of high-resolution structures produced from these pipelines, little evidence has been provided to justify such deviations from a standard cryo-ET workflow. So, we aimed to make clear just how impactful hotspots are on downstream data quality. The easiest possible method for montage tilt series collection is to acquire overlapping images with disregard for the accumulating dose. We tested exactly this on the thin leading edges of fibroblasts by manually arranging acquisition spots with a 10-20% overlap in both the X and Y dimensions (**Fig. 1a-c**). As expected, even with the smallest possible electron beam that still avoids image cutoffs and Fresnel rings, the beam outside of the imaged area nevertheless overlaps with the neighboring tiles, resulting in dose hotspots (**Supplementary Fig. 2**). Indeed, typical doses used for tomography (∼140 e^-^/Å^2^ for individual tiles), result in bubbling, especially near the centers of montages where the beam overlap is most severe (**Supplementary Figs. 3-5**). By contrast, however, merely decreasing the dose to ∼80 e^-^/Å^2^ largely avoids obvious signs of damage in all tested acquisition schemes (1×4, 2×2, 3×3, 4×1) (**Supplementary Figs. 3-5 and Supplementary Videos 1-3**). The resulting tilt series (**Supplementary Video 4**) and reconstructed montage tomograms (**Fig. 1f and Supplementary Video 5**) are generally seamless and retain sufficient detail. One can easily distinguish protein species, such as microtubules (**Fig. 1f**), filamentous actin, as well as cofilin-decorated actin^11^ (**Figure 2a-c and Supplementary Video 6)**. Using a 3×3 montage, we were even able to image a complete yeast cell. The cell wall and intracellular membranous compartments are easily identifiable and all ribosome positions can be annotated throughout the entire cell (**Figure 2d and Supplementary Video 7**). Thus, simply reducing the dose used for imaging produces high-quality montage tomograms by avoiding large-scale damage despite a high total electron dose in the hotspots.

**Fig. 2:**
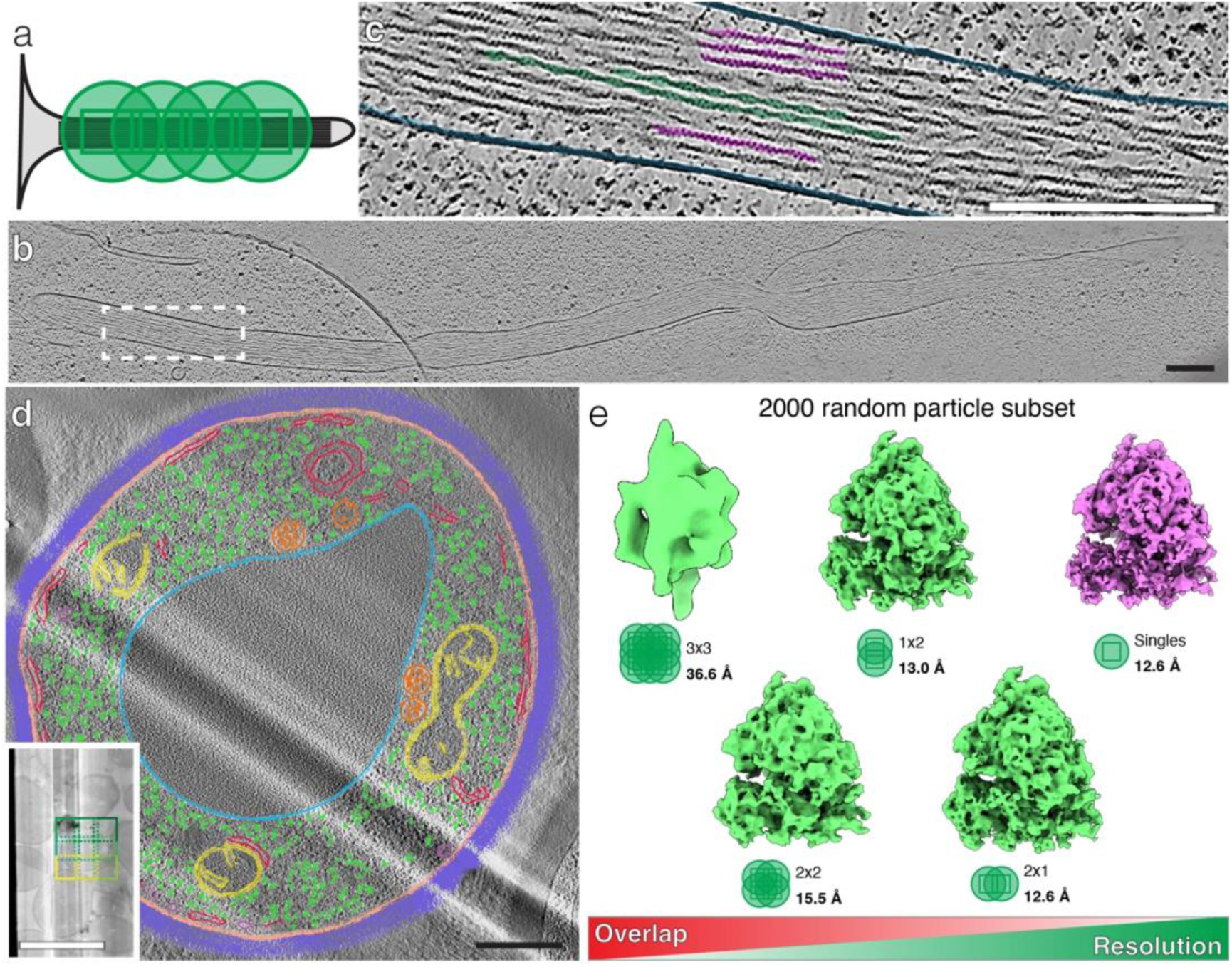
Products of the workflow. **a,** Cartoon representation of a 4×1 tile set being imaged along the long axis of a filopodium. **b,** A 7.2 nm-thick slice of a denoised, long, 4×1 montage tomogram of a filopodium from a 10T1/2 cell. Scale bar, 200 nm. **c,** Zoomed-in view of the spot in the white rectangle in **b**. Undecorated filamentous actin (magenta) and cofilin-decorated actin (green) are distinguishable constituents of the protrusion enclosed by the plasma membrane (blue). Scale bar, 200 nm. **d,** A segmented 3 nm-thick slice of a stitched, low-pass-filtered, 3×3 montage tomogram of an entire yeast cell. The locations of the constituent tilt series are denoted by the green and yellow squares on the lamella overview inset. The cell wall (violet), plasma membrane (light-orange), mitochondria (yellow), ribosomes (green), endoplasmic reticulum (red), vesicles (magenta), multivesicular bodies (dark-orange), and a vacuole (blue) are segmented. Scale bars, 500 nm, 5 µm (inset). **e,** Density maps from averaging a random subset of 2000 particles from various imaging modalities. In general, the resolution is inversely proportional to the degree of overlap, with a dramatic reduction in resolution for the 3×3 montage condition.

To more precisely measure the information retained or lost by accumulating dose unevenly during imaging, we performed subtomogram averaging on ribosomes from fibroblasts following different imaging schemes. Since the degree of beam overlap depends on the montage dimensions (**Supplementary Figs. 3-5**) we compared typical single tomograms with “long” (2×1; least overlap), “tall” (1×2), and square (2×2 and 3×3; most overlap) montages (**Supplementary Fig. 6**). Interestingly, 3D classification showed a similar proportion of good particles under each imaging regime with no obvious trend to observe (**Supplementary Fig. 7a**). Moreover, the good particles were evenly distributed across the tiles (**Supplementary Fig. 7b**), likely because at high tilts almost the entirety of the imaged area is exposed to more than one beam (**Supplementary Figs. 3,4**), leaving nearly no unaffected regions. As expected, by comparing reconstructions from single tomograms with an equal number of particles from different montage dimensions we found that there was an inverse correlation between the degree of overlap and the achievable resolution (**Fig. 2e**). The drop in resolution was modest for most montage dimensions tested, including small squares (2x2), but there was a marked decrease in resolution for the larger, 3x3 square montages (**Fig. 2e**) which have more severe dose accumulation towards their centers (**Supplementary Fig. 7c**). Thus, montages of large areas will apparently not be suitable for subsequent subtomogram averaging. However, when smaller montage tomograms are used, which already cover an area up to ∼2 µm^2^ in our setup, remarkable resolutions can still be achieved.

Fortunately, though dependent on beam size, tilt scheme, etc., even larger montages (4 tiles to a side and beyond) generally have comparable beam damage to a 3x3 montage, since additional tiles lie so far apart from one another (**Supplementary Fig. 7c**). To confirm this, as well as to further maximize the coverage of our montage tomograms, we attempted to collect large tile sets with lower magnifications but the same average electron dose. First, we experimented with low-magnification imaging on focused ion beam (FIB)-milled lamellae from epithelial cells grown on micropatterned EM grids (**Supplementary Fig. 8a,b**). After progressively stepping down in magnification (**Supplementary Fig. 9**), we settled on ∼6.5 Å/pixel because this reaches a large FOV while still preserving the visibility of small cytoplasmic components like actin filaments and ribosomes (**Supplementary Fig. 10 and Supplementary Video 8**). By acquiring large 4x5 montages at this magnification, we were able to cover areas over 80 µm^2^, which represents almost a complete lamella in a single tomogram (**Supplementary Fig. 8c,d and Supplementary Video 9**).

Finally, expanding our FOV through montage tomography would prove even more useful by applying these strategies to tissue where greater length scales and biological context can be probed. With muscle tissue as an example, previous implementations of cryo-ET on isolated myofibrils revealed unprecedented molecular detail of thin and thick filament structures^1,12,13^. However, individual tomograms of isolated sarcomeres lack cellular context and miss organelles, myofibril arrangements, etc. To solve this, we applied montage tomography to high-pressure frozen skeletal muscle tissue after FIB milling using the “waffle method”^14^ (**Supplementary Fig. 11**). Single tomograms from the resulting lamellae showed sarcomeric components and the adjacent sarcoplasmic reticulum (**Supplementary Fig. 11f**). Montage tomograms, however, enlarge the FOV and show multiple adjacent full-length sarcomeres, the sarcoplasmic reticulum, and nearby mitochondria (**Supplementary Fig. 11g and Supplementary Video 10**). Interestingly, even intermediate filaments are visible at the z-disc, supporting previous reports that they anchor the z-disc to the sarcolemma^15^.

Altogether, we have presented a streamlined workflow for montage cryo-ET using conventional image acquisition and MontageMaker for subsequent tile stitching. The workflow is straightforward and easily employable by any group interested in obtaining montage tomograms. For instance, MontageMaker requires minimal user input, can be used on data derived from any imaging software, and does so in batch mode (**Supplementary Fig. 1**), an important feature given the need for data processing steps to keep up with the ever-increasing speed of tomographic data collection^16–18^. Furthermore, we have shown that negative effects from foregoing equal dose distribution during imaging are mitigated by simply reducing the total electron dose, as evidenced by the lack of visible damage (**Supplementary Figs. 3-5**) and the striking resolution obtainable from smaller montages (**Fig. 2e**). For larger montages, though, the beam damage is too strong and does not allow for high-resolution reconstructions. However, this is paid off by a very large FOV, giving insights into the molecular architecture of entire cells and tissues (**Fig. 2d and Supplementary Figure 11**).

Since our presented montage cryo-ET workflow is easy to implement, we believe that it will open the door for many scientists in the field to add this technique to their portfolio. Montage cryo-ET is key to bridge the gap between high-resolution cryo-EM and volume imaging, and will thus play a critical role in elucidating the molecular mechanisms governing cell biology.

## Methods

### Cell culture and maintenance

10T1/2 fibroblasts and A-431 epithelial cells were purchased from LGC Group and maintained as prescribed by the supplier. Specifically, both cell types were cultured in DMEM (Gibco) containing GlutaMAX, 4.5 g/L D-glucose, and pyruvate, supplemented with 10% fetal bovine serum and 1% penicillin/streptomycin. Cells were maintained in T-75 flasks at 37°C and 5% CO_2_ and split every 2-3 days at somewhere between 60-90% confluency using 0.25% trypsin-EDTA (Gibco) for cell detachment.

*Saccharomyces cerevisiae* yeast cells were grown overnight at 30°C in YPD (1% yeast extract, 2% bactopeptone, 2% dextrose). Cultures were diluted the next morning in YPD to an optical density (OD) of ∼0.2 and were incubated for 4 additional hours at 30°C. Just prior to plunge-freezing, cells were diluted to 0.5 OD.

### Cryo-ET sample preparation and vitrification

For 10T1/2 cells, two types of EM grids were used. Both were from QUANTIFOIL and had a SiO_2_ film and a gold mesh (200). For initial testing, we used R 2/2 grids but used R 2/1s with an additional 2 nm of carbon for subtomogram averaging experiments. The grids were glow-discharged in a Quorum GloQube with 15 mA for 90 seconds and UV-sterilized for ∼10 minutes in a cell culture hood. Before cell seeding, grids were coated with 5 µg/mL fibronectin (Sigma-Aldrich) in 2 mL of PBS -/- in a 35 mm, glass-bottom MatTek dish for 30-60 minutes at room temperature. Then, grids were rinsed once in 2 mL PBS -/- per dish and 180,000 cells (∼20,000/cm^2^) were plated in 2 mL DMEM. 2-4 hours later, grids were plunge-frozen into liquid ethane using a Leica GP2 and the following settings: 37°C, 90% humidity, and one-sided sensor blotting for 2-5 seconds.

For A-431 experiments, grids (R 1/4 or R 2/2) were first glow discharged as above. Then, grids were micropatterned using a PRIMO 2 module (Alvéole) attached to a Zeiss Axio Observer Z1 widefield microscope. Here, grids were first coated in 0.01% PLL (Sigma-Aldrich; one 0.5 mL drop for 4 grids) for ∼1 hour at room temperature. Grids were then rinsed ∼4 times in Milli-Q H_2_O, dabbed dry with tissue paper, and placed in small drops (∼15 µL per grid) of 100 mg/mL PEG-SVA (Laysan Bio Inc.) in 100 mM pH 8.4 HEPES buffer for ∼1 hour at room temperature. Grids were again rinsed ∼4 times in Milli-Q H_2_O, then placed in a 35 mm MatTek dish containing HEPES buffer to await patterning. Patterning occurred on small (2-4 µL) drops of PLPP liquid and 20 µm circles were etched into the center of each grid square using a UV dose of 900 mJ/mm^2^. After patterning, grids were immediately rinsed in Milli-Q H_2_O and stored in a MatTek dish containing PBS in the dark at 4°C. Usually, grids were used the same day they were patterned. Grids were coated in the dark with 50 ug/mL FITC-fibronectin (Sigma-Aldrich) in PBS -/- for 30-45 minutes at room temperature. They were then submerged 3-4 times in PBS -/- to wash away excess fibronectin, and subsequently submerged in 1 mL DMEM (containing the supplements described above) in a 35 mm glass-bottom MatTek dish. 180,000-270,000 (∼20-30,000/cm^2^) cells in 1 additional mL of DMEM were added on top of the grids and briefly stirred to ensure even distribution. 2-3 hours later, grids were plunge-frozen as above after blotting for 5-7 seconds. Despite the thin ice, small cell size, and similar freezing conditions to the 10T1/2 fibroblasts, complete vitrification of A-431 cells proved difficult in our hands. This was particularly notable in large vacuole-like organelles (**Supplementary Fig. 8d**), suggestive of those being nucleation sites for hexagonal ice formation. In any case, this apparently had a limited impact on the low-resolution ultrastructural features we visualized here (**Supplementary Figs. 8-10**).

Yeast cells were diluted to 0.5 OD immediately prior to plunge-freezing. They were also frozen in the Leica GP2 with the following settings: 13°C, 80% humidity, and one-sided sensor blotting for 10 seconds.

Prior to tilt series collection, A-431 and yeast cells were FIB milled in an Aquilos 2 cryo-DualBeam FIB/SEM system (ThermoFisher Scientific) to a nominal lamella thickness of ∼100 nm.

### Fluorescence microscopy

When fluorescence microscopy was performed on EM grids, 10T1/2 cells were handled as above except, instead of plunge-freezing, cells were fixed in 1-2 mL of a 4% PFA (Electron Microscopy Sciences)/4% sucrose mixture in PBS for 10 minutes at room temperature and rinsed 3 times with 1-2 mL PBS. Then, CellMask Deep Red Actin Tracking Stain (Invitrogen; 1:500 dilution in PBS) was added for 90 minutes at room temperature. Grids were rinsed twice in 1-2 mL PBS and once in an equivalent volume of Milli-Q H_2_O before being mounted using a drop of ProLong Glass Antifade Mountant with NucBlue (Invitrogen) under a 12 mm #1.5 German glass coverslip (Electron Microscopy Sciences). For staining on glass, 10T1/2 cells were grown, fixed, and stained as above except they were first seeded onto 12 mm coverslips in 24-well plates, and these were mounted directly onto a glass slide after staining. Fluorescence imaging of cells and FITC-fibronectin was performed on a Zeiss LSM 800 with an Airyscan detector for fluorescence detection and an electronically switchable illumination and detection (ESID) module for brightfield imaging.

### Tilt series collection

Tilt series were acquired on a 300 kV Titan Krios (ThermoFisher Scientific), using two different hardware setups. For initial testing datasets on 10T1/2 (**Figs. 1f, 2b,c, and Supplementary Figs. 3-5**) and A-431 cells (**Supplementary Figs. 9,10**), images were collected on a non-fringe-free illumination (FFI) microscope equipped with a K3 direct electron detector (Gatan) and a zero-loss BioQuantum K3 energy filter (Gatan) set to a slit width of 20 kV. Other low-magnification A-431 montages (**Supplementary Fig. 8**), all yeast, skeletal muscle tissue, and subtomogram averaging data were collected on an FFI microscope with a Falcon 4i camera and a SelectrisX energy filter (both ThermoFisher Scientific). The SelectrisX energy filter was set to a slit width of 10kV. We saw no difference in tilt series collection or stitching between the two setups and most cartoons depicting imaging parameters use the rectangular K3 dimensions other than those in subtomogram averaging-related figures which use the square Falcon 4i dimensions. In either case, acquisition locations were set by manually moving “exposure” sites (as named in the software) in Tomography 5 (ThermoFisher Scientific) to where each tile had approximately a 10-20% overlap with its neighbors in both X (parallel to the tilt axis) and Y (perpendicular to the tilt axis).

For initial testing on 10T1/2 cells, tilt series were collected at 26kx magnification (3.58 Å/pixel) and a defocus of -8 µm. A bidirectional tilt scheme was used with a range of -51° to 51° and a tilt step of 3° or a range of -50° to 50° and a tilt step of 2°. Dosage varied from 50-140 e^-^/Å^2^ for each tilt series/tile. For subtomogram averaging of ribosomes, 10T1/2 cells were imaged with a magnification of 53kx (2.398 Å/pixel), a defocus range of -2 to -5.5 µm, a tilt range of -52° to 52° in a dose symmetric tilt scheme, a tilt step of 2°, and a dose of 80 e^-^/Å^2^. A-431 cells were imaged in the same manner except a series of magnifications were used including 8.7k x (11.27 Å/pixel on the K3), 11kx (8.68 Å/pixel on the K3), 15kx (6.66 Å/pixel on the K3), and 19.5kx (6.42 Å/pixel on the Falcon 4i) with one of the tilt ranges and step sizes listed above, a defocus of -8 µm, and a dose of 80 e^-^/Å^2^. Yeast cells were imaged on the Falcon 4i with a magnification of 33kx (3.77 Å/pixel), a tilt range of -51 to 51°, a tilt step of 3°, a defocus of -8 µm, and a total dose of 80 e^-^/Å^2^ per tile.

### Tilt series stitching and reconstruction

MontageMaker was written with assistance from ChatGPT versions 3.5 and 4^19^. It is written in Python and uses the tkinter^20^ package for its graphical user interface. MontageMaker uses the newstack command from IMOD^10^ to split tilt series into their constituent tilt images and renames the files according to the expectations of its core stitching script, a slightly modified version of that made by Yang et al^7^. Tile merging itself is done with the blendmont command of IMOD.

Overlapping tilt series were merged using MontageMaker according to the basic procedure shown in **Fig. 1e** and **Supplementary Fig. 1**. In short, directories containing tilt series were imported into the software and tiles were arranged according to the serpentine order expected by the stitching script^7^ in the “Tile Selection” tab. The instructions for tile rearrangement are also accessible from within the software. The queue, if merging multiple montages, is then set. Next, tilt ranges and increments are specified in the “Stitching Parameters” tab and the tilt series are split into their individual tilt images and renamed to fit with the expected inputs of the stitching script. Most other parameters (camera size and approximate tile overlap) are determined automatically by the software. Finally, after entering the X and Y tile dimensions, all montages in the queue are stitched sequentially. A summary text file that includes the calculated FOV of the final montage as well as the montage tilt series itself are then available for viewing from the “Results” tab. Detailed instructions can be found on a GitHub link that will be made prior to publication.

Montage tilt series were reconstructed using weighted-back projection in the Etomo module in IMOD versions 4.10.51 or 4.11.10^10^.

### Subtomogram averaging

If subtomogram averaging was to be performed, tilt series were collected at 53kx (2.398 Å/pixel) in EER format (180 frames per tilt image). Although the images could be stitched into montages, we performed all further processing, picking, etc. on the individual montage tiles for simplicity (**Supplementary Fig. 6**). The frames were then motion-corrected and their CTFs estimated in Relion 5^21^. Tilt series alignment and reconstruction were also performed using the IMOD wrapper in Relion 5. 4-fold binned tomograms were used for particle picking in crYOLO^22^. Tomograms were then sorted by image modality for a final tally of 64 single tomograms, 7 3x3 montages (63 tomograms), 16 2x2 montages (64 tomograms), 17 1x2 montages (34 tomograms), and 16 2x1 montages (32 tomograms). Particles were then extracted at bin 2 (4.796 Å/pixel) in Relion. Each group underwent 3D classification with 3 classes and a ribosome (PDB ID 7CPU^23^) low-pass filtered to 100 Å was used as an initial reference. In each case other than 3x3 montages there was one good class and two junk classes. We were unable to isolate a clear good class from 3x3 montage tomograms, and therefore went forward with the one that most resembled a ribosome. The good classes contained 3889 (singles), 14139 (3x3s), 5373 (2x2s), 3266 (1x2s), and 2118 (2x1s) particles. A subset of 2000 random particles were made from all groups for appropriate comparison with the others. These classes underwent further 3D refinement with the map from the final classification iteration used as the initial reference. Then, we performed post processing with a tight mask and a user-assigned B-factor of -200 resulting in reconstructions of 12.6 (singles), 36.6 (3x3s), 15.5 (2x2s), 13.0 (1x2s), and 12.6 Å (2x1s) in resolution according to gold-standard FSC criteria. Local resolution was determined on the 3D refinement results using Relion and colored on the refinement map using ChimeraX^24^. The entire subtomogram averaging workflow is shown in **Supplementary Fig. 6**.

It is worth noting that by processing individual tiles, the particles that exist in the overlapping regions are duplicated. For the purposes of this study, we did not exclude them from the analysis as they were not true duplicate particles (same signal and same noise), but were actually distinct images of the same protein. It is possible that if one were to seek high resolution from this method that repeated copies of these particles would need to be deleted before classification and/or refinement.

### Tomogram post-processing and segmentation

For better visualization, tomograms were either low-pass filtered (60-180 Å) using EMAN2^25^ or denoised in Dragonfly version 2022.2^26^, as specified in the respective figure legends. For Dragonfly denoising, we used a regressive neural network trained on a “synthetic cytoplasm” simulated using cryo-TomoSim^27^. Briefly, multiple noisy synthetic tomograms were generated with varied simulation parameters and then trained to regress to a noiseless “prior” that was generated from the same tomographic model. Tomogram segmentation was performed with a neural network trained in the Segmentation Wizard of Dragonfly and the results were manually cleaned, also in Dragonfly.

### Cryo-ET of skeletal muscle tissue

Soleus muscles were dissected from 16-day-old BALB/c mice as described previously^1^ and homogenized to achieve dissociated muscle fibers of ∼50 µm in diameter and lengths between 100 µm and 1 mm. Homogenization was performed in an Omni THQ Digital Tissue Homogenizer using 3-second rounds at 5000 rpm in rigor buffer^1^ at 4°C (usually 2-3 rounds). Before freezing, fibers were stained with CellMask Deep Red Actin Tracking Stain (Invitrogen, 1:1000 dilution in rigor buffer) for 60 minutes at 4°C. Excess stain was removed via 3 rounds of centrifugation at 2500 rpm for 5 minutes at 4°C in rigor buffer.

Muscle tissue was high-pressure frozen in a Leica ICE using a modified version of the “waffle method”^14^. EM grids (R 2/2, 200 mesh, gold, UltrAuFoil; QUANTIFOIL) were glow discharged as above and flipped such that the support film faced downward (**Supplementary Fig. 11a**). In this orientation, 5 µL of sample were added to the grid and excess buffer was manually blotted from beneath with filter paper. Grids were then placed onto the flat face of a 1-hexadecene-coated carrier and moved into a middle plate slot before a 50 µm gold spacer was placed on top. 3 µL of rigor buffer supplemented with 10% dextran was placed on top of the muscle fibers before assembling the sandwich and freezing.

Grids were clipped foil-side up (**Supplementary Fig. 11a**) and loaded into an Aquilos 2 cryo-DualBeam FIB/SEM (ThermoFisher Scientific) on a 35° shuttle. Milling sites were chosen based on fluorescence correlation using a METEOR integrated fluorescence microscope (Delmic). Images of actin fluorescence were manually aligned with SEM grid overviews in the MAPS software (ThermoFisher Scientific; **Supplementary Fig. 11b-d**). Milling was performed using the waffle method^14^, except it began at 54° and proceeded in ∼10° steps until the final milling angle of 8°. Lamellae were polished to a nominal thickness of ∼200 nm.

Tilt series collection was performed as above on the Falcon 4i camera at 33kx (3.77 Å/pixel), a tilt range of -51 to 51°, a tilt step of 3°, a defocus of -8 µm, and a total dose of 80 e^-^/Å^2^. Montages were stitched as above with MontageMaker, and tomograms were reconstructed using 5 iterations of SIRT in Etomo. Tomogram contrast was enhanced using low-pass filtering in EMAN2^25^.

## Data Availability

All relevant data are available from the corresponding author upon reasonable request. All tomograms shown in the Supplementary Videos and all subtomogram averages will be deposited in the Electron Microscopy Data Bank (EMDB) and made available prior to publication.

## Code Availability

MontageMaker source code will be deposited on GitHub and made publicly available prior to publication.

## Acknowledgements

We thank O. Hofnagel, D. Prumbaum, and K. Kubat for electron microscopy technical support and Thorsten Wagner for help with particle picking in crYOLO. Mice were provided by the Leibniz Research Centre for Working Environment and Human Factors (IfADo) in Dortmund, Germany. This work was supported by funds from the Max Planck Society (to S.R.) and the European Research Council under the European Union’s Horizon 2020 Programme (ERC-2019-SyG, grant no. 856118 (to S.R.). R.H. and M.B.S. are supported by a postdoctoral fellowship from the Alexander von Humboldt foundation.

## Author contributions

R.H. and S.R. designed the study. R.H. cultured and plunge froze all mammalian cell samples and performed all fluorescence microscopy experiments. M.B.S performed yeast cell culture and supported R.H. with their plunge freezing and FIB milling. A.P. prepared mouse muscle tissue samples including dissection, high-pressure freezing, and FIB milling. R.H. designed and wrote MontageMaker and performed all tilt series collection, montage tile stitching, tomogram reconstruction, and segmentation. G.R. and R.H performed subtomogram averaging. R.H. wrote the initial manuscript and R.H., M.B.S, and S.R. revised it with contributions from all authors.

## Declaration of interests

The authors declare no competing interests.

---

**Supplementary Figure 1:**
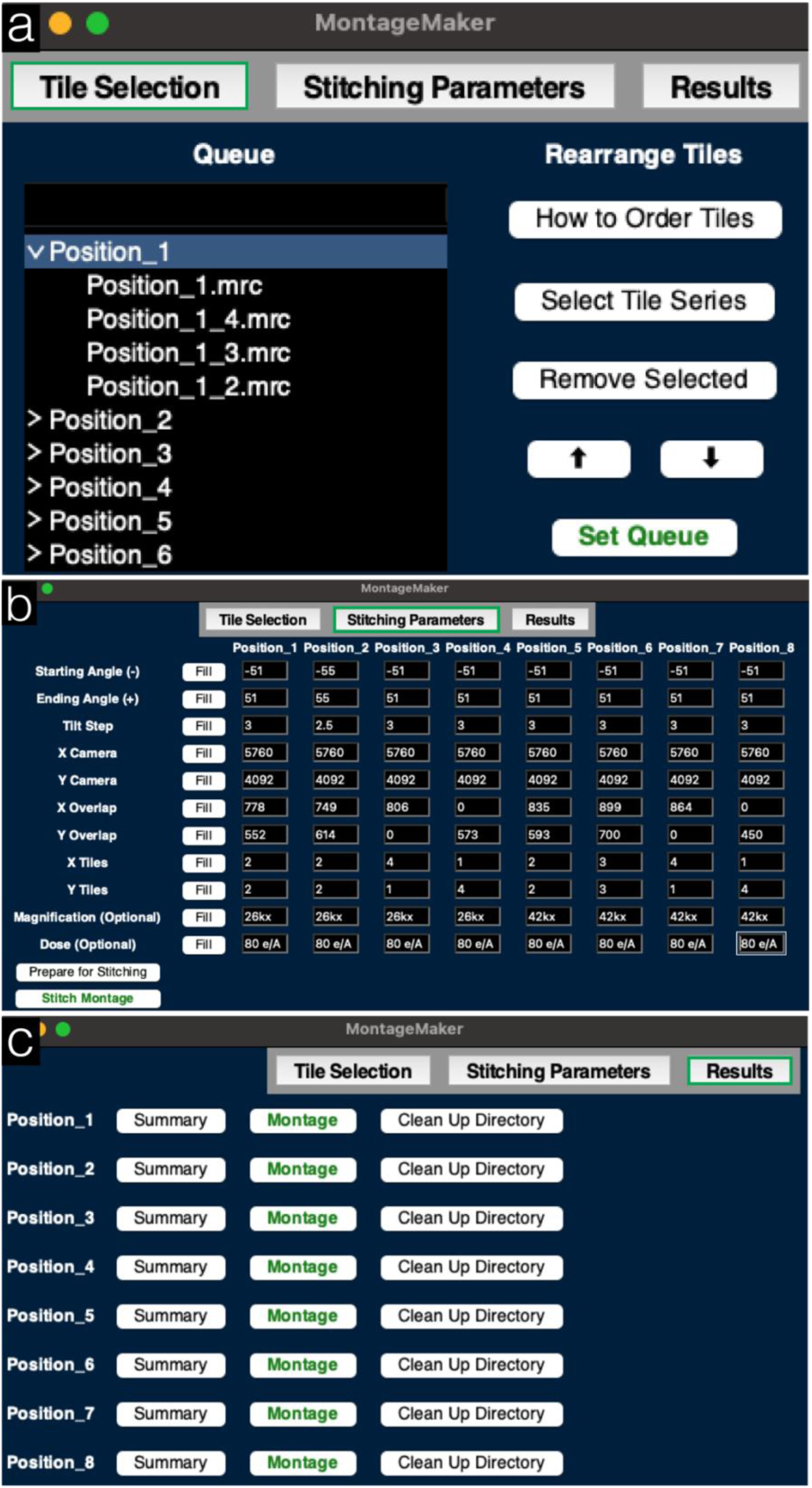
MontageMaker workflow. **a-c**, The three main windows in the MontageMaker software: “Tile Selection” (**a**), “Stitching Parameters” (**b**), and “Results” (**c**).

**Supplementary Figure 2:**
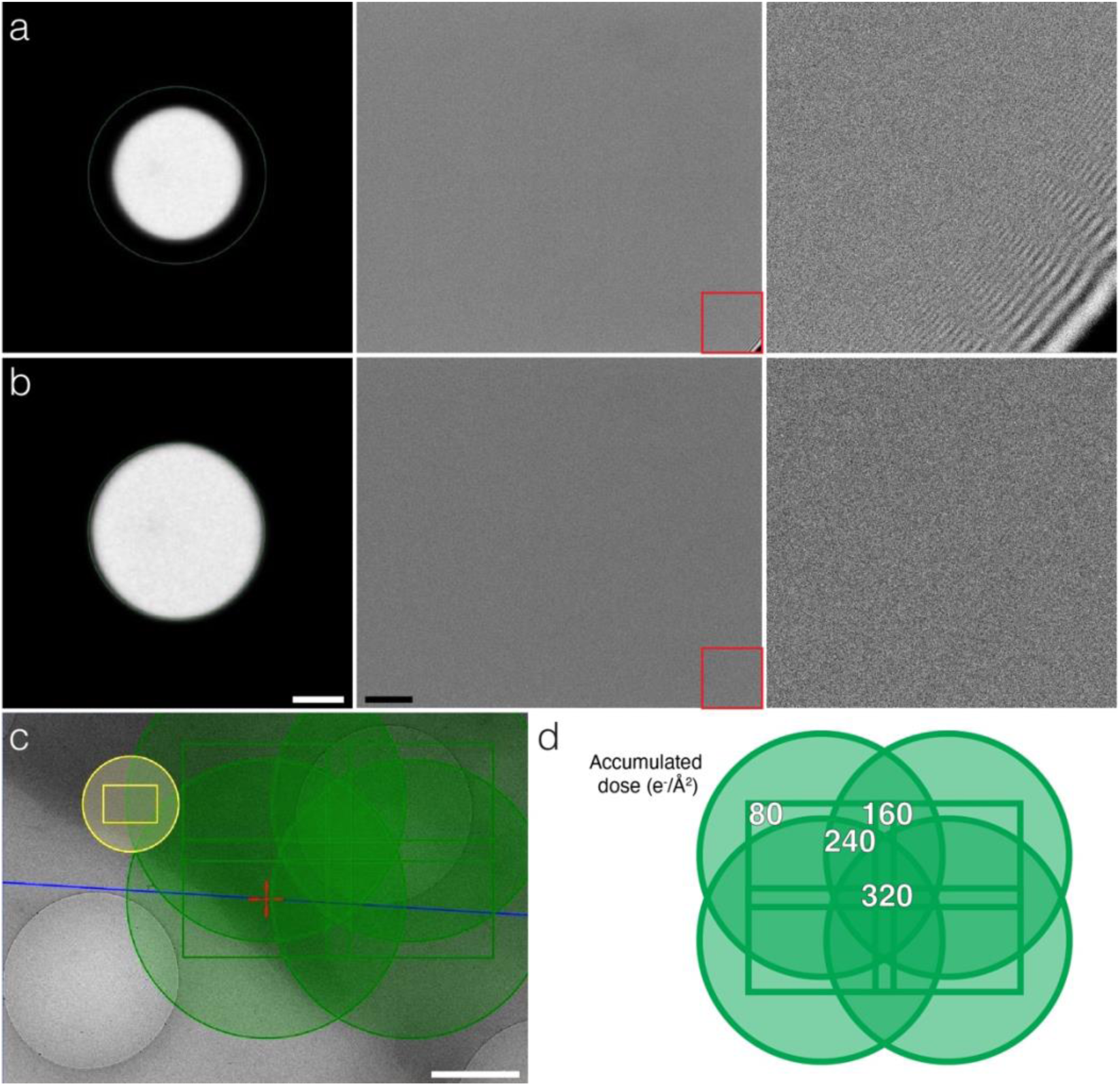
Electron beam dimensions for initial testing. **a,b,** The electron beam was made as small as possible while still avoiding image cutoffs and fringes. While a 2.6 µm beam results in artifacts (**a**), a 3.45 µm (**b**) beam avoids them). The left column shows the fluorescent screen display on the TEM user interface, the middle column an image on the camera (K3 camera on Gatan’s Digital Micrograph software), and the right column has a zoomed-in view of the middle insets shown in red. Scale bars, left panels, 1 µm; middle panels, 200 nm **c,** Screenshot displaying overlapping tilt series being collected using Tomography 5. Scale bar, 1 µm. **d,** Despite a small beam, overlap is inevitable and results in dose hotspots in the montage tomogram. Shown here is an example 2x2 tile set with an 80 e^-^/Å^2^ total dose per individual tilt series, a 3.45 µm beam diameter, and 26 kx magnification on a rectangular K3 camera. The values shown indicate the total dose across the final montage with the given overlap (here 15%).

**Supplementary Figure 3:**
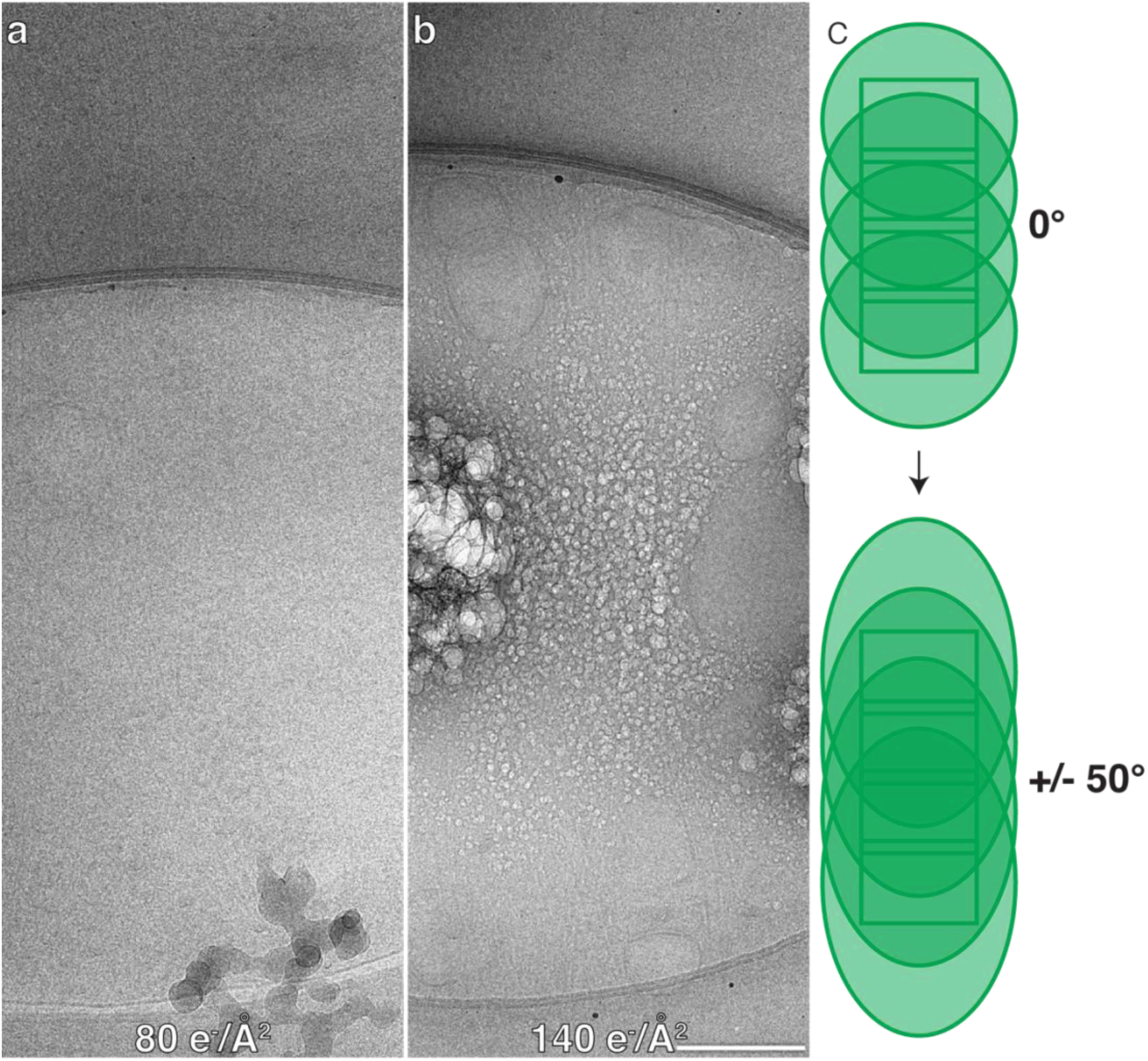
Beam damage is reduced on montages with low total dose. **a,b,** Final stitched tilt image (50°) of a 1x4 tilt series from 10T1/2 fibroblast lamellipodia imaged with either 80 e^-^/Å^2^ (**a**) or 140 e^-^/Å^2^ (**b**). Here, the tilt axis is running from left to right perpendicular to the long axis of the montage. Obvious radiation damage is visible only on the 140 e^-^/Å^2^ montage. Images were acquired on a rectangular K3 camera. Scale bar, 200 nm. **c,** Cartoon diagrams showing the increased overlap at higher tilt angles for a 1x4 montage where the initial overlap in Y is 15% of the camera height. Beam diameters at 0° are 3.45 µm.

**Supplementary Figure 4:**
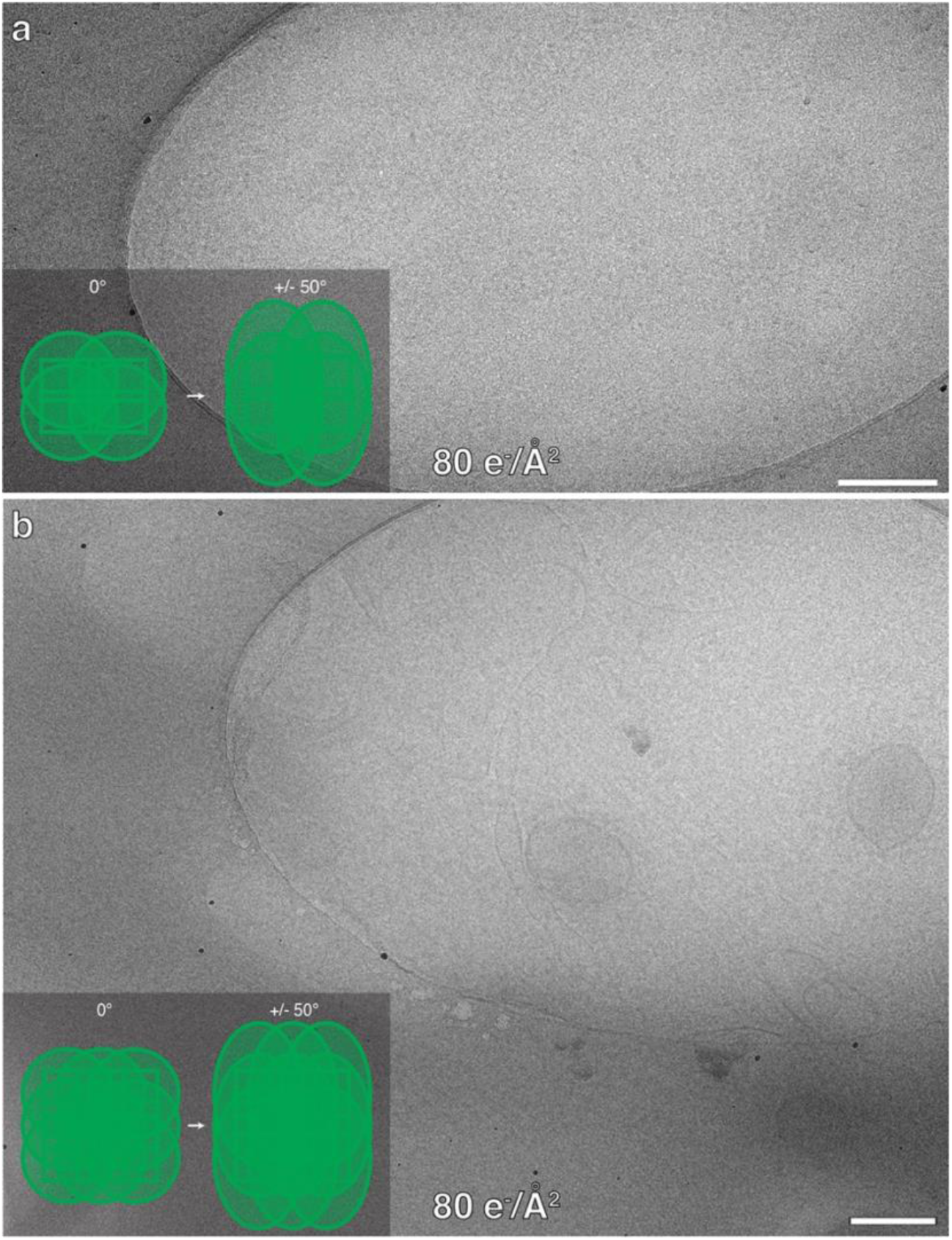
Beam damage is minimal in square montages with low total dose. **a,** Final stitched tilt image (50°) of a 2x2 tilt series from a 10T1/2 fibroblast lamellipodium with 80 e^-^/Å^2^. Minimal beam damage is visible in the center of the image where the beam overlap between adjacent tiles is most severe. The inset shows a cartoon diagram showing the overlap at initial and final tilt angles for a 2x2 montage where the initial overlap in X and Y is 15% of the camera length/height. Beam diameters at 0° are 3.45 µm. Images were acquired on a rectangular K3 camera. Scale bar, 200 nm. **b,** Same as **a**, but for a 3x3 montage. Here, the grid’s film is slightly damaged, but all cellular structures remain intact. Images were acquired on a square Falcon 4i camera. Scale bar, 200 nm.

**Supplementary Figure 5:**
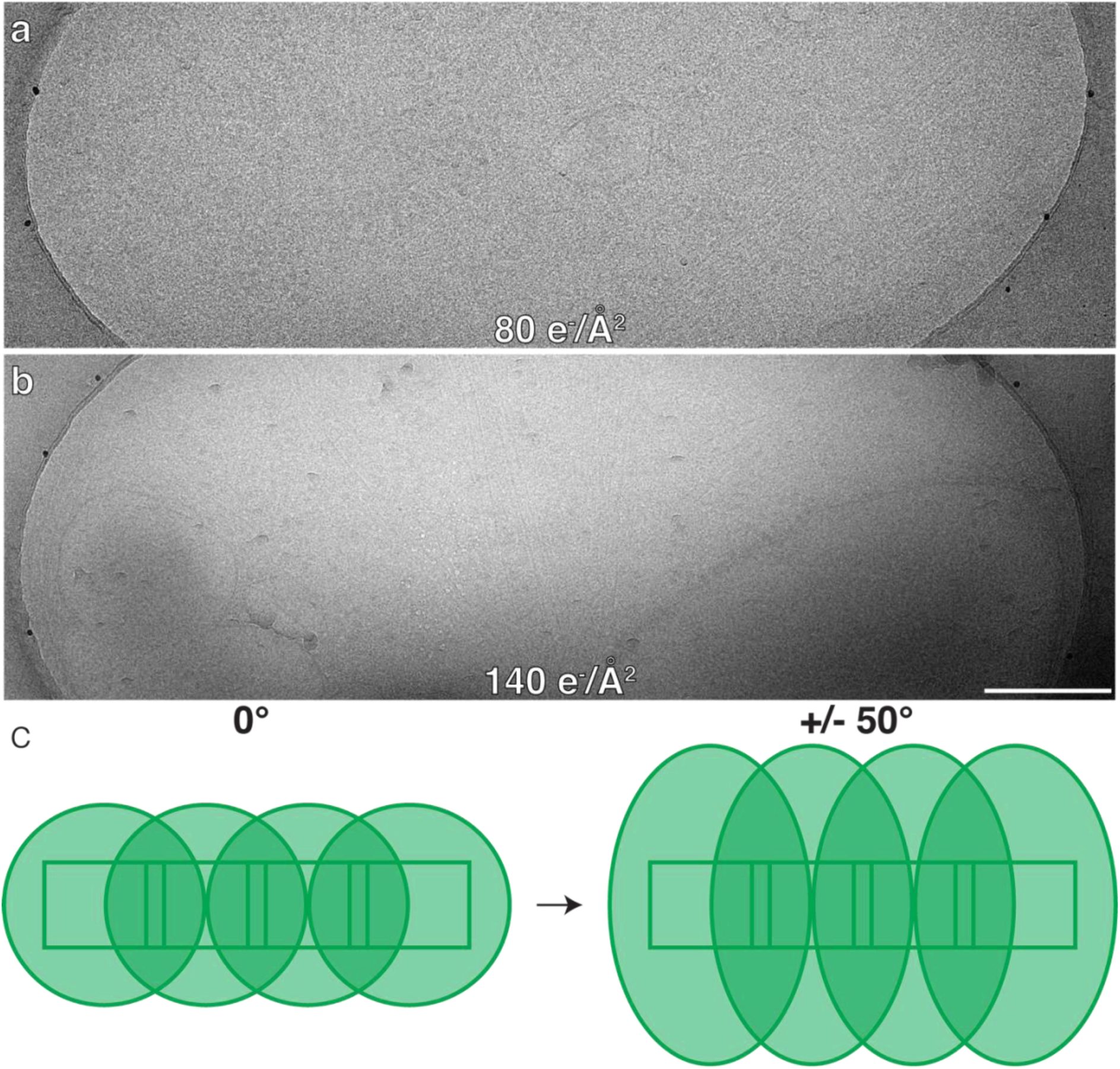
Beam damage is minimal in unidimensional montages whose long axis is parallel to the tilt axis with low total dose. **a,b,** Final stitched tilt image (50°) of a 4x1 tilt series from 10T1/2 fibroblast lamellipodia imaged with either 80 e^-^/Å^2^ (**a**) or 140 e^-^/Å^2^ (**b**). Here, the tilt axis is running from left to right along the long axis of the montage. Only minimal beam damage is visible, even at a total dose 140 e^-^/Å^2^. Images were acquired on a rectangular K3 camera. Scale bar, 200 nm. **c,** Cartoon diagram showing the overlap at initial and final tilt angles for a 4x1 montage where the initial overlap in X is 15% of the camera length. Beam diameters at 0° are 3.45 µm.

**Supplementary Figure 6:**
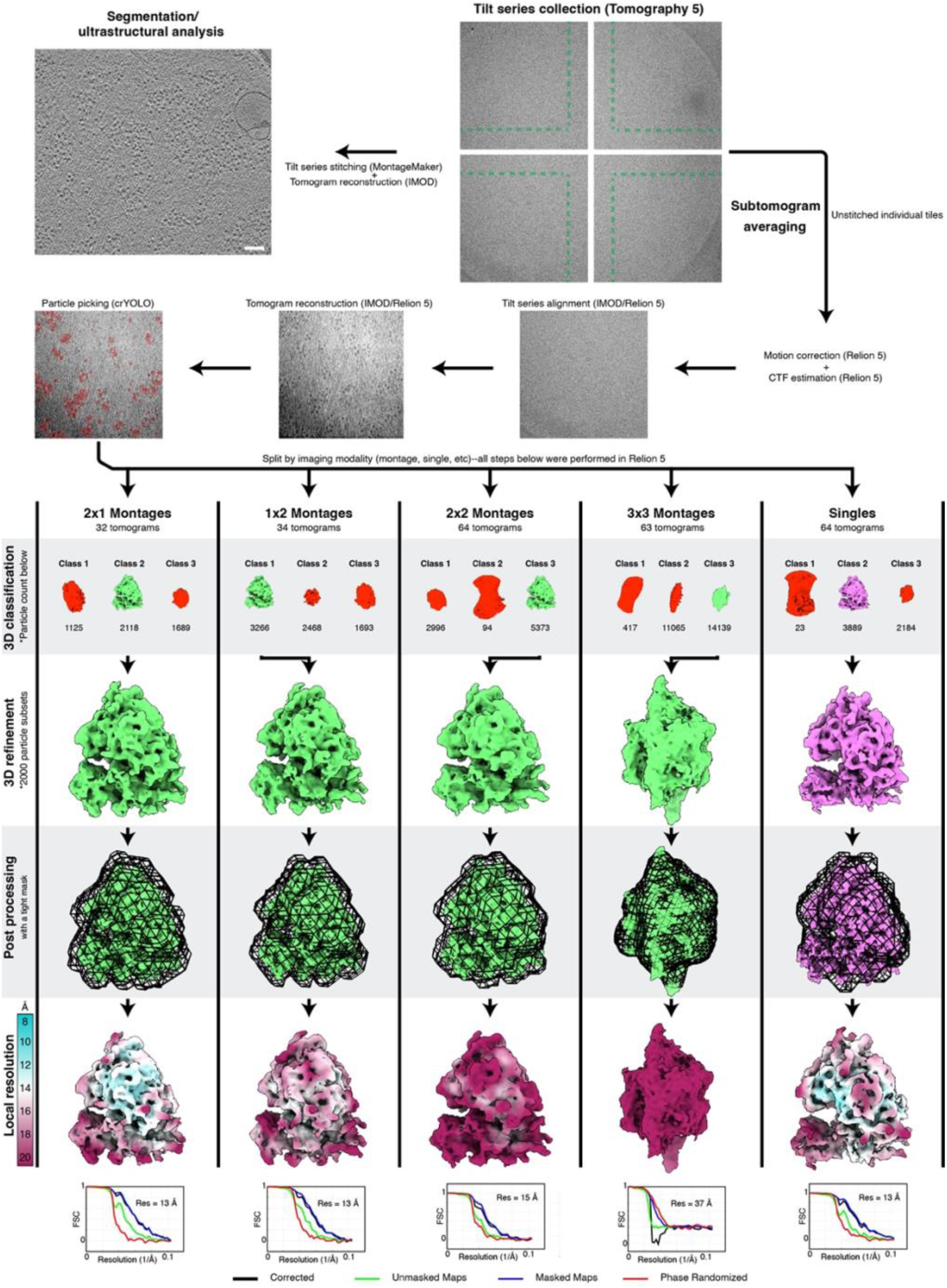
Subtomogram averaging workflow. Although tilt series are stitched into montages before tomogram reconstruction for segmentation and ultrastructural analysis, we processed unstitched individual tiles during subtomogram averaging. The green dashed line in the tiles indicates the approximate region of overlap with the other tiles. Motion correction and CTF estimation were performed in Relion 5 and ribosomes were picked using crYOLO. All further classification and refinements were done using Relion 5. First, each imaging modality was separated and was processed on its own. 3D classification then split the particles into 3 groups. The best class from each group was further refined. For comparison with the other groups, subsets of 2000 random particles were used. Finally, a tight mask was applied to each map and post-processing was performed to sharpen the map and arrive at the final gold standard resolution. Scale bar, 100 nm.

**Supplementary Figure 7:**
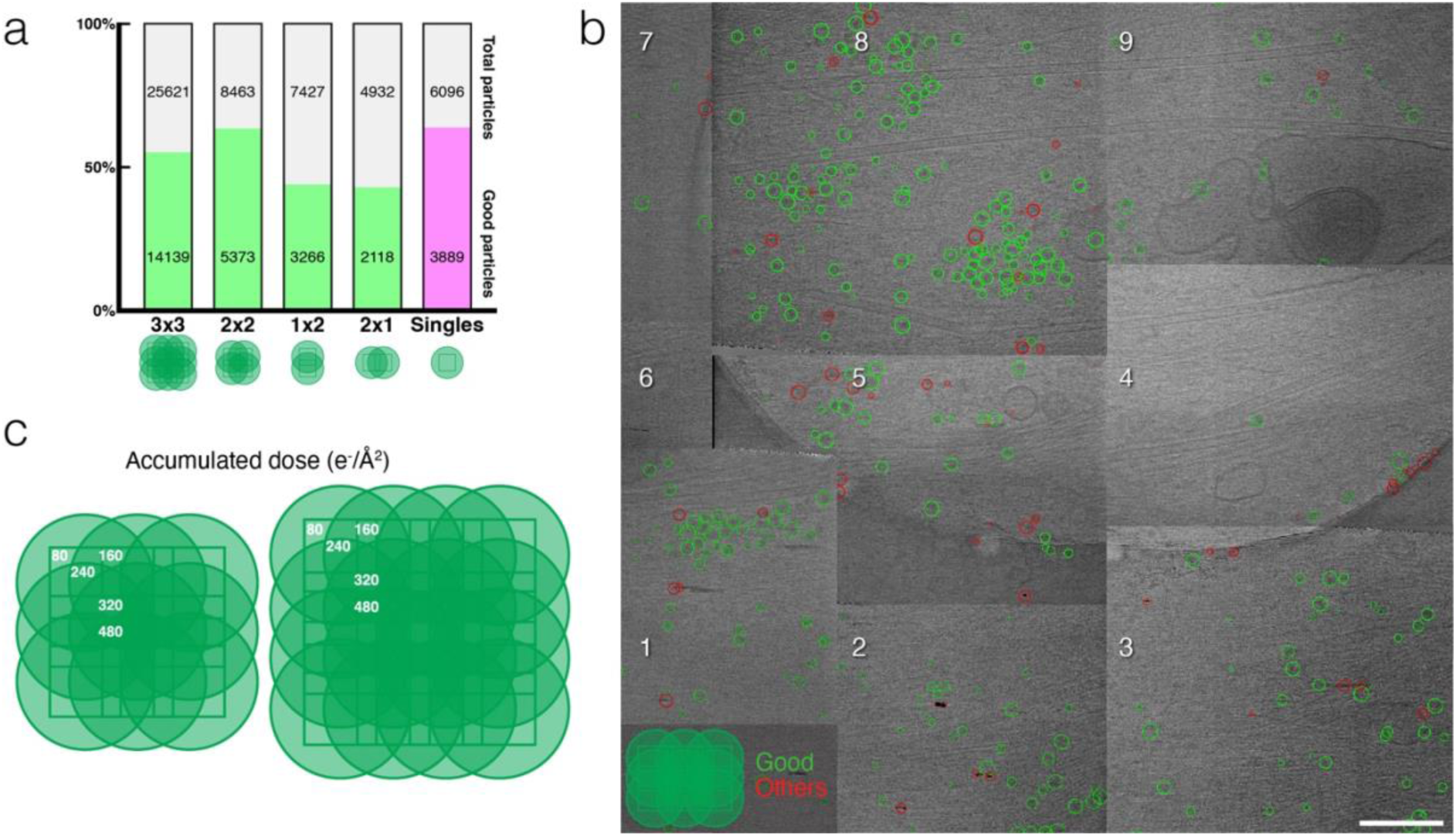
Ribosome classification results and dose accumulation comparisons across montage sizes. **a,** Counts for the particles that were well classified compared to the overall number of particles for each imaging modality. **b,** Example 3x3 montage tiles (unstitched and overlaid manually) with ribosomes from the good (green) and other (red) classes displayed. No clear trend is visible for particle distributions from either class and most poorly classified particles (red) are not ribosomes. Scale bar, 250 nm. **c,** Example 3x3 and 4x4 montage tile sets showing the total accumulated dose across different hot spots given a dose of 80 e^-^/Å^2^ for each tile.

**Supplementary Figure 8:**
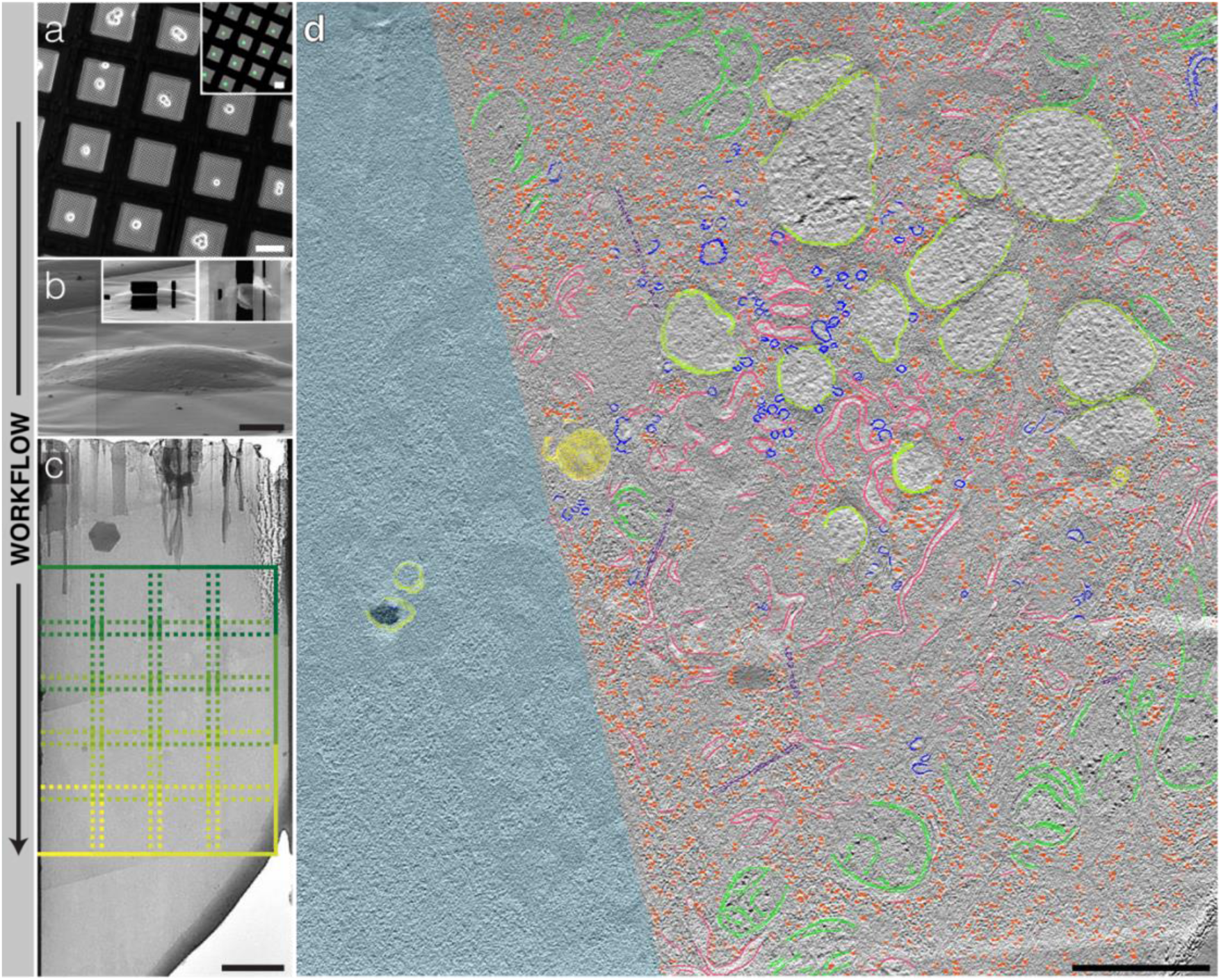
Whole-lamella montage tomography of a A-431 cell. **a,** A-431 cells seeded onto a micropatterned EM grid. The pattern consists of 20 µm circles etched into the center of each grid square and coated with FITC-fibronectin (inset, green). Scale bars, 50 µm. **b,** An A-431 cell before (main) and after (insets) FIB-milling. The lamella shown is ∼600 nm thick prior to fine polishing. Scale bar, 5 µm. **c,** Image locations of the 4x5 montage tomogram in **d** which covers nearly the entire lamella and 82 µm^2^ of cellular area. Scale bar, 2 µm. **d,** 12.8 nm-thick slice of a montage tomogram acquired at 19.5kx on a Falcon 4i camera (6.42 Å/pixel) and low-pass-filtered. Microtubules (violet), endoplasmic reticulum (pink), vesicles (blue), ribosomes (orange), mitochondria (green), multilamellar bodies (yellow), and vacuole-like organelles (yellow-green) are segmented. The nucleus is covered in light blue. After reconstruction and trimming, ∼50 µm^2^ of area are displayed here. Scale bar, 1 µm.

**Supplementary Figure 9:**
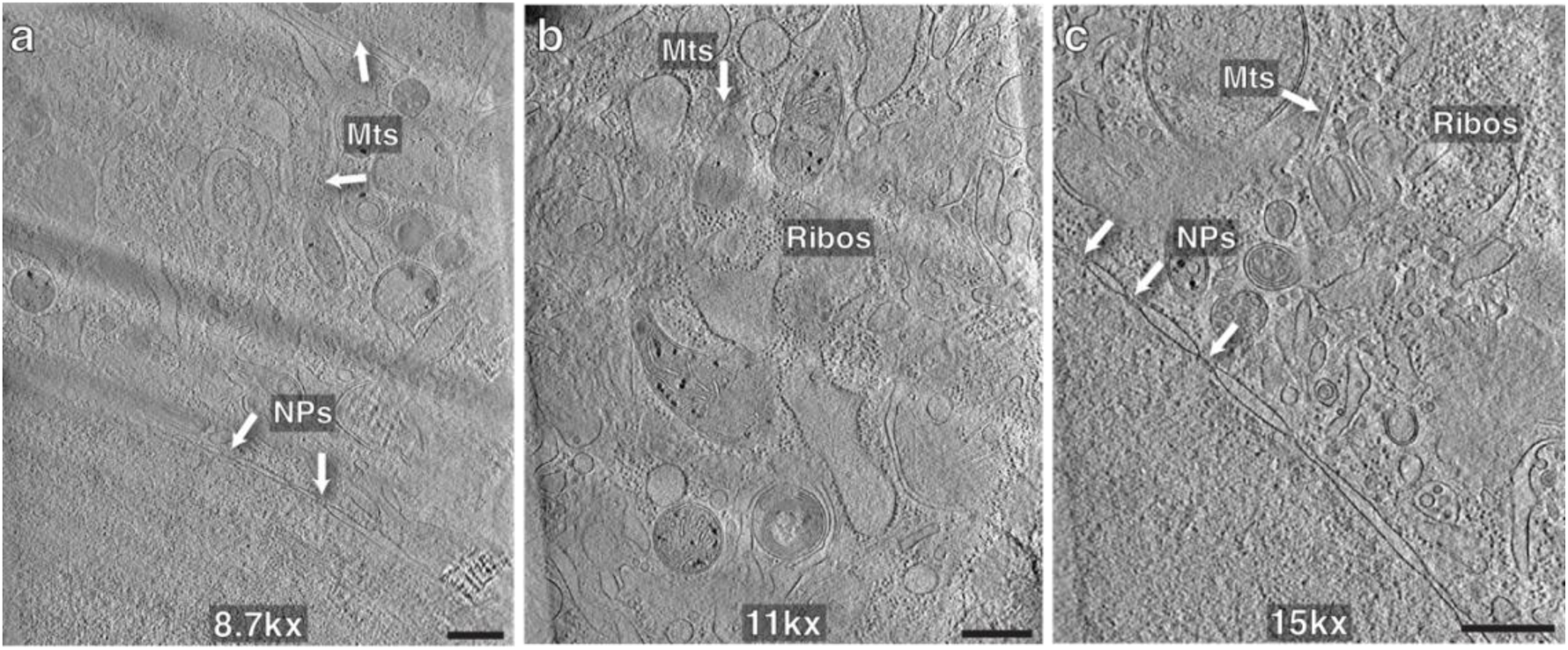
Magnifications tested for low-magnification montage tomography. **a-c,** Slices from individual low-pass-filtered tomograms acquired from A-431 cells at 8.7kx (11.27 Å/pixel) (**a**), 11kx (8.68 Å/pixel) (**b**), or 15kx (6.66 Å/pixel) (**c**) on our K3 camera. Slices are 22.5 nm (**a**), 17.4 nm (**b**), or 26.7 nm (**c**) thick. Example microtubules (Mts), ribosomes (Ribos), and nuclear pores (NPs) are labeled. Scale bars, 500 nm.

**Supplementary Figure 10:**
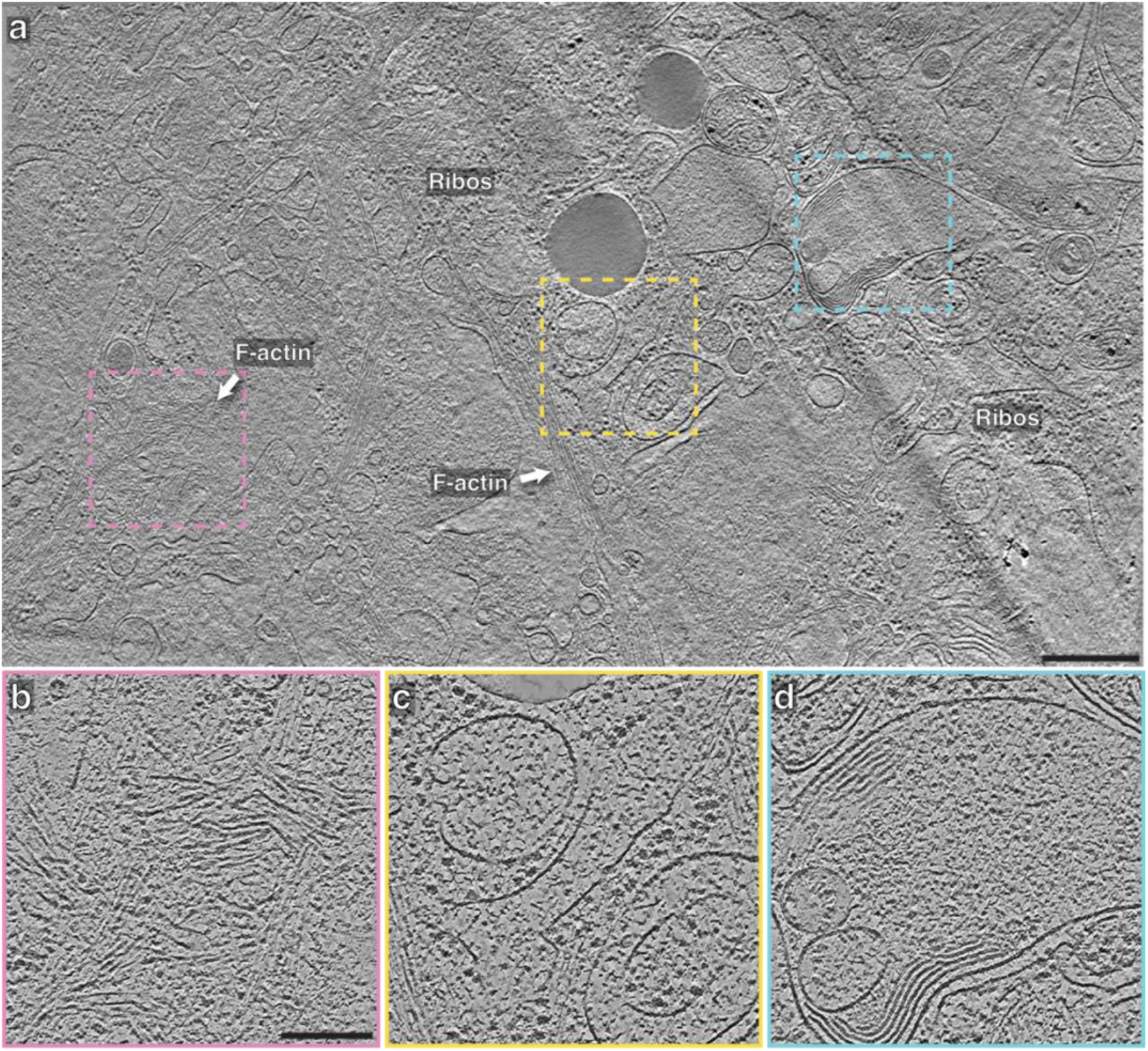
Low-magnification montage tomogram with small cytoplasmic components visible. **a,** 2.7 nm-thick slice of a 2x2 low-pass-filtered montage tomogram acquired on a lamella of an A-431 cell. Despite the low magnification (15kx, 6.66 Å/pixel), small features are visible such as actin filament bundles (F-actin) and ribosomes (Ribos). Scale bar, 500 nm **b-d,** Denoised and zoomed-in views of the pink (**b**), yellow (**c**), and blue (**d**) squares in **a**. Scale bar, 200 nm.

**Supplementary Figure 11:**
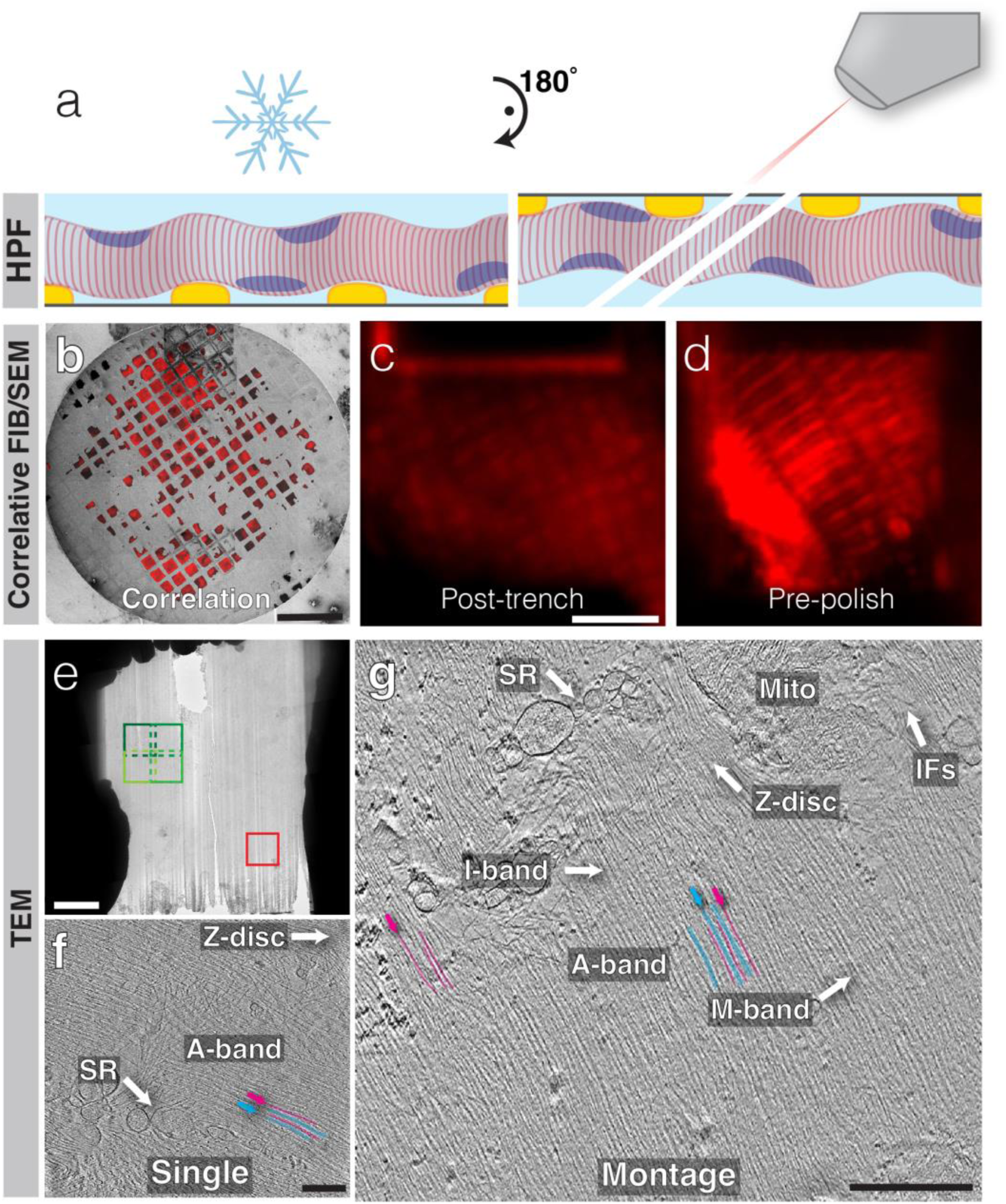
Pipeline for montage cryo-ET of skeletal muscle tissue. **a,** Cartoon representation of the grid and sample orientation during high-pressure freezing and during FIB milling (rotated 180° with respect to the high-pressure freezing setup). **b,** Representative fluorescence overlay atop an SEM grid overview. Actin (CellMask Deep Red Actin Tracking Stain) is red. Scale bar, 500 µm. **c,d,** In-FIB/SEM fluorescence images of a lamella after trench milling (**c**) and a few steps before polishing (**d**). Scale bar, 10 µm. **e,** TEM lamella overview. The locations of the single tomogram (red) and montage (green) in **f** and **g**, respectively, are shown. Scale bar, 5 µm. **f,** 9.1 nm-thick slice through a single low-pass-filtered tomogram showing sarcomeric components and sarcoplasmic reticulum (SR). Magenta and cyan lines/arrows point out example thin and thick filaments, respectively. Scale bar, 250 nm. **g,** 9.1 nm-thick slice of a low-pass-filtered montage tomogram showing multiple sarcomeres, in addition to sarcoplasmic reticulum (SR), intermediate filaments (IFs), and a mitochondrion (Mito). As in **f**, thin and thick filament examples are denoted by the magenta and cyan lines/arrows, respectively. Scale bar, 500 nm.

**Supplementary Video 1: 1x4 montage tomogram from a 10T1/2 lamellipodium.** This tomogram is comprised of four tilt series that overlap perpendicular to the tilt axis and was collected at 26kx on a K3 camera. This is the tomogram from Supplementary Figure 3a.

**Supplementary Video 2: 3x3 montage tomogram from a 10T1/2 lamellipodium.** This tomogram is comprised of nine tilt series arranged in a 3x3 pattern and was collected at 53kx on a Falcon4i camera. This is the tomogram from Supplementary Figure 4b.

**Supplementary Video 3: 4x1 montage tomogram from a 10T1/2 lamellipodium.** This tomogram is comprised of four tilt series that overlap parallel to the tilt axis and was collected at 26kx on a K3 camera. This is the tomogram from Supplementary Figure 5a.

**Supplementary Video 4: Stitched montage tilt series are typically seamless.** This montage tilt series is near the leading edge of a 10T1/2 cell and is comprised of four tilt series arranged in a 2x2 square pattern collected at 26kx on a K3 camera. This tilt series was reconstructed into the tomogram shown in Figure 1f and Supplementary Video 5.

**Supplementary Video 5: 2x2 montage tomogram from a 10T1/2 lamellipodium.** This tomogram is comprised of four tilt series arranged in a 2x2 pattern and was collected at 26kx on a K3 camera. This is the same tomogram as is shown in Figure 1f and Supplementary Video 4.

**Supplementary Video 6: Denoised 4x1 montage tomogram of a filopodium from a 10T1/2 cell.** This tomogram is comprised of four overlapping tilt series collected at 26kx on a K3 camera that run parallel to the long axis of the filopodium. This is the same tomogram as in Figure 2b,c.

**Supplementary Video 7: 3x3 montage tomogram of an entire yeast cell.** Nine tilt series were arranged in a 3x3 pattern to cover a whole yeast cell and parts of its neighbors. Tilt series were collected at 33kx on a Falcon 4i camera. This is the same tomogram as in Figure 2d.

**Supplementary Video 8: 2x2 low-magnification montage tomogram from an A-431 cell lamella.** Tilt series were collected at a low (15kx/6.66 Å/pixel) magnification and were arranged in a 2x2 pattern. Tilt series were collected on a K3 camera. This is the same tomogram as in Supplementary Figure 10.

**Supplementary Video 9: 4x5 low-magnification montage tomogram of nearly a complete A-431 cell lamella.** Tilt series were collected at a low (19.5kx/6.42 Å/pixel) magnification and were arranged in a 4x5 pattern. Tilt series were collected on a Falcon 4i camera. This is the same tomogram as in Supplementary Figure 8d.

**Supplementary Video 10: 2x2 montage tomogram from mouse soleus muscle tissue.** Tilt series were collected at 33kx on a Falcon 4i camera and were arranged in a 2x2 pattern. This is the same tomogram as in Supplementary Figure 11g.

## Notes

### Competing Interest Statement

The authors have declared no competing interest.

